# *miR-100* and *miR-125b* Contribute to Enhanced 3D Growth and Invasiveness and can be Functionally Transferred to Silence Target Genes in Recipient Cells

**DOI:** 10.1101/2024.01.16.575716

**Authors:** Hannah M. Nelson, Shimian Qu, Liyu Huang, Muhammad Shameer, Kevin C. Corn, Sydney N. Chapman, Nicole L. Luthcke, Sara A. Schuster, Lauren A. Turnbull, Lucas L. Guy, Xiao Liu, Kasey C. Vickers, Qi Liu, Jeffrey L. Franklin, Alissa M. Weaver, Marjan Rafat, Robert J. Coffey, James G. Patton

## Abstract

Extracellular communication via the transfer of vesicles and nanoparticles is now recognized to play an important role in tumor microenvironment interactions. Cancer cells upregulate and secrete abundant levels of *miR-100* and *miR-125b* that can alter gene expression by both cell- and non-cell-autonomous mechanisms. We previously showed that these miRNAs activate Wnt signaling in colorectal cancer (CRC) through noncanonical pairing with 5 negative regulators of Wnt signaling. To identify additional targets of *miR-100* and *miR-125b*, we used bioinformatic approaches comparing multiple CRC cell lines, including knockout lines lacking one or both of these miRNAs. From an initial list of 96 potential mRNA targets, we tested 15 targets with 8 showing significant downregulation in the presence of *miR-100* and *miR-125b*. Among these, Cingulin (CGN) and Protein tyrosine phosphatase receptor type-R (PTPRR) are downregulated in multiple cancers, consistent with regulation by increased levels of *miR-100* and *miR-125b.* We also show that increased cellular levels of *miR-100* and *miR-125b* enhance 3D growth and invasiveness in CRC and glioblastoma cell lines. Lastly, we demonstrate that extracellular transfer of *miR-100* and *miR-125b* can silence both reporter and endogenous mRNA targets in recipient cells and also increase the invasiveness of recipient spheroid colonies when grown under 3D conditions in type I collagen.

## 1. Introduction

MicroRNAs (miRNAs) are small (22-23nt) RNAs that base pair with target mRNAs to regulate gene expression by decreasing translation and promoting RNA degradation (He and Hannon, 2004, Krol et al., 2010). In cancer, miRNA expression patterns are frequently altered, contributing to all aspects of tumor progression from proliferation to metastasis (Peng and Croce, 2016). We and others have focused on the role of *miR-100* and *miR-125b* in cancer (Cha et al., 2015, Qin et al., 2015, Peng et al., 2021, Liu et al., 2022, Lu et al., 2017, Zhang et al., 2021). These miRNAs are encoded in the third intron of *MIR100HG*, a 3kb lncRNA that is transcribed from the large 330kb MIR100HG locus (Emmrich et al., 2014). Increased expression of *miR-100, miR-125b* and *MIR100HG* can promote cetuximab (anti-EGFR) resistance in colorectal cancer (CRC) cells, drive progression of pancreatic cancer, and promote the epithelial to mesenchymal transition (EMT)(Lu et al., 2017, Liu et al., 2022, Ottaviani et al., 2018). These effects are driven in part by regulating the expression of target mRNAs including mTOR (Cha et al., 2015), negative regulators of Wnt signaling (Lu et al., 2017), Cystic Fibrosis Transmembrane Regulator (CFTR) (Zhang et al., 2021), Cingulin (CGN) (Zhang et al., 2021), and checkpoint kinases (CHK) (Ottaviani et al., 2018). Beyond these mRNAs, TargetScan predicts a large number of additional potential targets for *miR-100* (59 targets) and *miR-125b* (931 targets)(https://www.targetscan.org/vert_80/). Base pairing between miRNAs and target 3’-UTRs is typically imperfect, with canonical targets displaying extensive pairing between nucleotides 2-8 of the miRNA (the “seed” sequence) and variable complementarity throughout the rest of the miRNA (Bartel, 2009). However, even within the seed sequence, the extent of base pairing between miRNAs and their mRNA targets is highly variable making prediction of mRNA targets a challenge (Li et al., 2008, Riolo et al., 2021). Indeed, all 5 of the negative regulators of Wnt (DKK1, DKK3, ZNRF3, RNF43, and APC2) that we previously found to be targeted by *miR-100* and *miR-125b* are noncanonical targets, raising the question as to whether additional targets might contribute to cetuximab resistance and altered 2D and 3D growth.

Extracellular vesicles (EVs) and nanoparticles participate in cell-cell communication through the transfer of RNA, protein, and lipids (Couch et al., 2021, Dixson et al., 2023). Among RNA cargo, transfer of miRNA is the most well characterized due to their small size and the ability to experimentally monitor silencing of target mRNAs in recipient cells using reporter constructs (O’Brien et al., 2020). We previously showed that *miR-100* and *miR-125b* are secreted from CRC cells with Transwell co-culture experiments supporting extracellular transfer of these miRNAs to recipient cells (Cha et al., 2015). However, since the recipient cells in those transfer experiments expressed endogenous *miR-100* and *miR-125b*, it remained a possibility that the observed silencing could be due to unexpected activation of the endogenous genes or other indirect effects (Gruner and McManus, 2021).

To identify additional targets of *miR-100* and *miR-125b* and to definitively demonstrate extracellular transfer, we created cell lines with CRISPR/Cas9-mediated knockouts of *miR-100, miR-125b,* or both. Through extensive bioinformatic analyses of RNAseq data, we identified and tested 15 candidate targets for *miR-100* and *miR-125b* with the two most statistically significant being Cingulin (CGN) and Protein Tyrosine Phosphatase Receptor Type-R (PTPRR). We also show that *miR-100* and *miR-125b* contribute to enhanced 3D growth and invasiveness of CRC and glioblastoma cells. Importantly, we demonstrate functional transfer of *miR-100* and *miR-125b* from donor to recipient cells leading to silencing of both reporter constructs and endogenous genes, as well as increased invasiveness of spheroid cultures grown in type I collagen.

## 2. Materials and Methods

### 2.1 Cell Culture

The CRC cell line HCA-7 was plated in 3D culture in type I collagen to generate CC cells (cystic colonies)(Lu et al., 2017). CC cells were then subjected to iterative selection in 2D (cetuximab resistant) and 3D (cetuximab sensitive) conditions until a resistant line (CC-CR) was generated that is capable of growth in 3D in the presence of cetuximab (Lu et al., 2017). The LN-18 glioblastoma line (ATCC CRL-2610) was a kind gift from the lab of Dr. Michael King at Vanderbilt. All cell lines were confirmed to be free of mycoplasma contamination (Venor^TM^GeM Mycoplasma Detection Kit, PCR-based, Sigma Aldrich Cat #MP0025). Cells were grown in Dulbecco’s Modified Eagle’s Medium (DMEM, Corning) supplemented with 10% fetal bovine serum (FBS, Corning), 1% L-glutamine (200 nM, Gibco), 1% MEM nonessential amino acids (Sigma Aldrich), and 1% penicillin-streptomycin (10,000 U/mL, Gibco) in 5% CO_2_ at 37°C. Cells were passaged a maximum of 8 times before discarding.

### 2.2 Generation of Knockout Cell Lines

We used CRISPR/Cas9 multiplex genome engineering (Kabadi et al., 2014) to delete *miR-100, miR-125b,* or both. Guide RNAS (gRNAs) flanking the targeted locus were designed using the Benchling CRISPR gRNA Design Tool (https://www.benchling.com/crispr). Oligonucleotides containing gRNA sequences were first cloned into sgRNA expression vectors (Addgene #53186-53189) and then multiple gRNA cassettes were cloned into lentiviral expression vectors (pLV hUbC-Cas9-T2A-GFP; Addgene, #53190) using Golden Gate cloning technology. Lentiviral stocks were prepared in 293T cells and used to transduce CC-CR cells. Single GFP^+^ cells were sorted into 96 cell plates and expanded. PCR amplification of genomic DNA from clonal GFP^+^ cells was performed using primers flanking the gRNA target sequences to identify deletion clones.

### 2.3 Extracellular Vesicle Isolation

Cells were plated at 9.0×10^6^ cells in T-175 flasks (Corning), with each EV collection using at least 3 flasks. Cells were grown to 80% confluency and washed 3 times with 1X PBS before media were replaced with DMEM lacking FBS and grown for 48 hours. Culture media were then collected and adherent cells were trypsinized and counted. Conditioned media were then centrifuged at increasing speeds and time: 1,000xg for 10 minutes (room temperature), 2,000xg for 20 minutes (4°C), and 10,000xg for 30 minutes (4°C). Crude EV pellets were obtained after the conditioned media was centrifuged for 17 hours at 100,000xg (4°C). EV pellets were then washed two times by resuspension in 1mL of 1X DPBS (Corning) and centrifugation at 100,000xg for 70 minutes. Final pellets were resuspended in 100µl 1X DPBS. Under these conditions, the EV pellets consist of a heterogenous mixture of EVs with some non-vesicular material (Jeppesen et al., 2019). This protocol was used to directly compare with earlier results examining export of *miR-100 and miR-125b* into CRC EVs (Cha et al., 2015). EVs were quantified by protein concentration (Pierce BCA Protein Assay) and by Nanoparticle Tracking Analysis (NTA, ZetaView Nanoparticle Tracking Analysis Instrument).

### 2.4 qRT-PCR

Total RNA from both whole cell and EVs was isolated using TRIzol (Life Technologies). Taqman small RNA assays (Life Technologies) were used to quantify miRNA levels. A total of 10ng of RNA were utilized for each RT reaction; 0.67 µl of cDNA was used in each 10µl qPCR reactions. qPCR was completed in either 96-well or 384-well plates using Biorad CFX96 or CFX384 instruments. Fold changes were calculated as previously described (Cha et al., 2015).

### 2.5 Transfer Assays

For Transwell co-culture assays, cells were plated in Transwell dishes with 0.4µm filters (Corning, 3460) with 50,000 cells in the donor well and 300,000 cells in the recipient well. Transwell co-cultures were grown for five days, changing media every 72 hours. Cells were washed 3 times with 2 mL of 1X PBS; RNA was then collected from either donor or recipient cells.

### 2.6 AGO2 Immunoprecipitation

For immunoprecipitation, anti-AGO2 (clone 11A9 antibody; MABE253, Sigma-Aldrich) or control IgG antibodies were used. Ten 150cm dishes of either CC or CC-CR cells were grown to 95% confluency. Cells were then scraped in ice cold PBS and centrifuged to collect cell pellets in ice cold lysis buffer (20mM Tris HCl pH 7.5, 150 mM KCl, 0.5% NP40, 2mM EDTA, 1mM NaF, 0.5mM DTT). After centrifugation of insoluble cell debris, cell lysates were incubated with Dynabeads. Prior to cell lysis, Dynabeads™ Protein G (ThermoFisher, 10003D) were conjugated with antibodies by incubating overnight with rotation at 4°C and washed two times with ice cold lysis buffer. Cell lysates were then added to the conjugated beads and incubated at 4°C for six hours. Beads were gently washed with fresh lysis buffer and proteins were eluted. For RNA isolation, beads were incubated with DNase and Proteinase K for 10 minutes at room temperature. Trizol was then added and incubated at room temp for 10 minutes and RNA was isolated.

### 2.7 Small RNA Sequencing

RNA libraries were generated using the NEXTFlex Small RNA Library Preparation Kits v3 (Perkin) with the following modifications: (1) 3′- and 5′-adaptors were diluted 1:8, (2) 3′-adaptor ligations were performed overnight in multiple steps –25°C for 2 h, 20°C for 4 h and 16°C overnight, (3) following cDNA synthesis and before barcoding PCR, step F protocol was followed, and (4) PCR amplification was 20 cycles. Following PCR amplification, individual libraries were size-selected (136–200 bp product) using Pippin Prep (Sage Sciences). Size-selected libraries were quantified using a Qubit Fluorometer. Libraries were checked for quality and sequenced using Illumina short-read technology. Libraries were pooled and paired-end sequencing (PE-150)(equimolar multiplexed libraries) was performed on the NovaSeq6000 platform by the VANTAGE core (Vanderbilt University). Demultiplexing and bioinformatic analyses were performed using the TIGER pipeline (Allen et al., 2018). Briefly, Cutadapt (v1.16) was used to trim 3′ adaptors and all reads with <16 nucleotides (nts) were removed (Martin, 2011). Quality control on both raw reads and adaptor-trimmed reads was performed using FastQC (v0.11.9)(www.bioinformatics.babraham.ac.uk/projects/fastqc). The adaptor-trimmed reads were mapped to the hg19 genome, with additional rRNA and tRNA reference sequences using Bowtie1 (v1.1.2), allowing only one mismatch (Langmead and Salzberg, 2012).

### 2.8 Total RNA Sequencing

Bulk RNA sequencing libraries were prepared using Universal RNAseq kits (Tecan). Libraries were cleaned (Zymo), checked for quality using the Agilent bioanalyzer, quantified (Qubit), and pooled based on equimolar concentrations. Pooled libraries were sequenced using Illumina short-read technology based on PE150 on the NovaSeq6000 (Illumina). After sequencing, samples (libraries) were demultiplexed and analyzed. Briefly, adapter sequences were removed using Cutadapt v2.10)(Martin, 2011). Quality control of both raw reads and adaptor-trimmed reads was performed using FastQC (v0.11.9). After adaptor trimming, reads were aligned to the Gencode GRCh38.p13 genome using STAR (v2.7.8a)(Dobin et al., 2013). FeatureCounts (v2.0.2)(Liao et al., 2014) was used to count the number of reads mapped to each gene. Heatmap3 (Zhao et al., 2014) was used for cluster analysis and visualization. Differential expression was analyzed by DESeq2 (v1.30.1)(Love et al., 2014). Significant differentially expressed genes were determined with fold change >2 or <0.5, and adjusted p values (padj) <0.05. Genome Ontology and KEGG pathway over-representation analyses were performed using the WebGestaltR package (NULL)(Wang et al., 2017). Gene set enrichment analysis (GSEA) was performed using GSEA package (v4.3.2)(Subramanian et al., 2005) on database v2022.1.Hs.

### 2.9 Gene Expression Analysis

The Gene Expression database of Normal and Tumor Tissues (GENT2) was used to compare levels of expression between 72 paired normal and tumor tissues utilizing the GPL570 platform. Statistical significance was determined using two-sample t-tests (GPL570 platform (HG-U133_Plus_2).

### 2.10 Transfection of Luciferase Plasmids

Cells were grown to approximately 50-60% confluency with transfections performed using TransIT®-2020 (Mirus, MIR5404). The Renilla luciferase construct (pClneo-RL, plasmid #115366, Addgene) was transfected at 0.5ug/mL and the firefly luciferase construct was transfected at 1ug/mL. DNA and TransIT-2020 were incubated for 30 minutes at room temperature in Opti-MEM (Gibco). Transfection complexes were diluted 1:10 in DMEM and added to cells.

### 2.11 Luciferase Reporter Assays

3’-UTRs from mRNAs of interests were cloned using RT/PCR into the pMIR-REPORT™ miRNA Expression Reporter Vector System (Firefly Luciferase) using NEBuilder HiFi DNA Assembly (New England BioLabs). PCR amplification primers are listed in **Table S1**. All plasmids were verified by sequencing (Genewiz). Transfected cells were cultured alone or in co-cultures using Transwells with 0.4μm membranes for 72 hours followed by washing with 1X PBS. Luciferase levels were then detected using the Dual-Glo® Luciferase Assay System (Promega). Changes in firefly luciferase levels were normalized to Renilla luciferase levels and compared to luminescence from the empty (no 3’UTR) vector.

### 2.12 Immunofluorescent Staining and Imaging of Cells

Cells were plated on coverslips and grown for three days after which coverslips were washed three times in 1x PBS and fixed in 4% paraformaldehyde for 30 minutes at room temperature. Samples were then washed and allowed to block for 1 hour at room temperature in blocking buffer (1X PBS with 0.001% Triton-X and 0.03% donkey serum). Primary antibody incubation (CGN, ab244406) was overnight at 4°C, followed by three 1X PBS wash steps, and incubation with secondary antibodies (Cy3 anti-Rabbit antibody and 488 conjugated phalloidin) for 2 hours at room temperature. Samples were washed three times with 1X PBS and mounted on glass slides using Vectashield with DAPI and imaged using a Zeiss LSM 880 microscope (Vanderbilt University, CISR). All images were analyzed using ImageJ (Aebi et al., 1986).

### 2.13 Spheroid Cultures

Cells (10,000) were plated in Nunclon Sphera low adherent plates (Thermo Scientific) and centrifuged at 1,000xg for 10 minutes at room temperature, as previously described (Carey et al., 2013). Cells were grown for three days before transfer into 2mg/ml Type I collagen (Advanced Biomatrix). Colonies were imaged using Muvicyte Live Cell imager for five days. Images were processed and analyzed using ImageJ(Aebi et al., 1986).

### 2.14 Overexpression of Cingulin

The Cingulin (CGN) coding region was amplified by RT/PCR from RNA isolated from CC cells using primers as shown in Table S1. PCR products were cloned into a Sleeping Beauty-based Tet-On vector (pSB829) using NEBuilder HiFi DNA assembly. The resulting construct was confirmed by sequencing. For transfections, 0.5 ug plasmid DNA (pCDNA3.1-SB100X) and 0.5 ug of overexpression vector were incubated with 2ug of Lipofectamine 2000 (ThermoFisher) in Opti-MEM (Gibco) for 30 minutes at room temperature. Transfection complexes were diluted in 1:10 in complete DMEM and added to LN-18 cells and incubated for 24 hours. Cells were then selected using 300 ug/mL of hygromycin. Cingulin expression was induced by adding 300ng/mL of doxycycline (DOX).

### 2.15 Protein Collection and Western Blotting

Proteins were collected using 1X-RIPA buffer (Life Technologies) and quantified using BCA assays (ThermoFisher). 40ug of total protein were loaded onto pre-cast SDS gels (4-20% Mini-PROTEAN TGX 50uL pre-cast gels; BIO-RAD). Separated proteins were transferred to nylon membranes using the Trans-Blot Turbo Transfer System (BIO-RAD). Membranes were blocked with Intercept Blocking Buffer (IBB) (LI-COR) for 1 hour at room temperature. Primary antibodies were incubated overnight in IBB at 4°C. Blots were washed three times in 1X TBS-T. Secondary antibodies were incubated for 1 hour at room temperature in IBB. Blots were then washed in 1X TBS-T three times. Blots were imaged using the Odyssey XF (LI-COR) and quantified using ImageJ.

## 3. Results

To facilitate identification of mRNA targets for *miR-100* and *miR-125b* and to develop cell lines to test for functional miRNA transfer, we used CRISPR/Cas9 technology to generate deletions of these two miRNAs within the MIR100HG locus (**Fig. 1A**). A single lentiviral vector expressing multiple gRNAs, Cas9, and GFP was created and lentiviruses were transduced into cetuximab resistant CC-CR cells (Lu et al., 2017). The gRNAs were designed to base pair with regions flanking *miR-100, miR-125b,* or both (**Fig. S1**). Targeting both *miR-100* and *miR-125b* also results in deletion of *let-7a* and the BLID open reading frame, but previous work did not support a role for these genes in cetuximab resistance (Lu et al., 2017). Individual GFP+ clones were sorted, targeted deletions were identified by PCR, and verified by sequencing. While CC-CR cells express high levels of *miR-100* and *miR-125b*, the parental, cetuximab sensitive CC cell line expresses ∼50-fold less of both miRNAs (Lu et al., 2017)(**Fig. 1B**). As expected, the knockout lines showed undetectable levels of *miR-100 or miR-125b* or both **(Fig. 1C)**.

**Figure 1.**
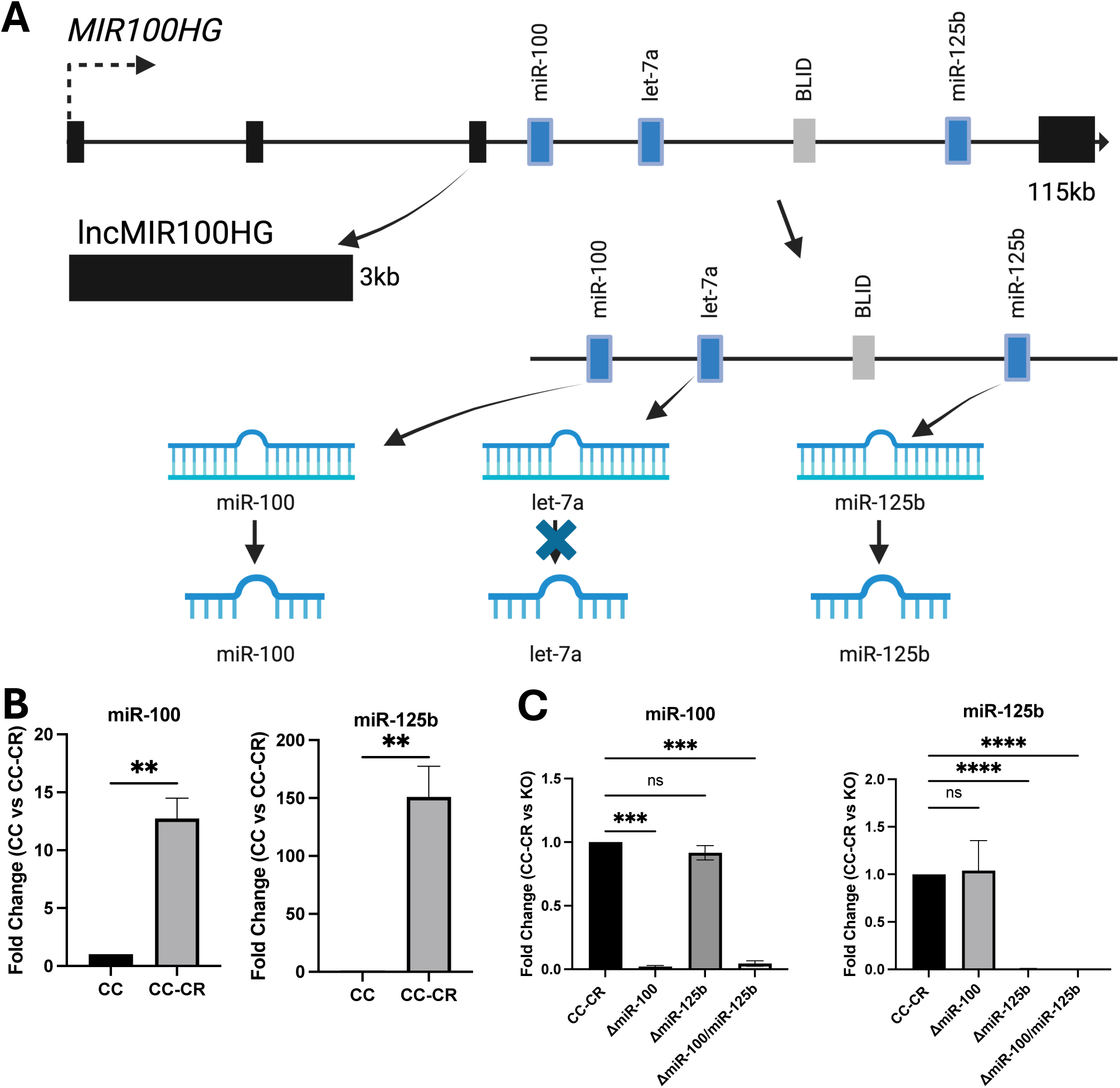
Expression levels of *miR-100* and *miR-125b* in CC, CC-CR and Δ*miR-100/miR-125b* cell lines. (A) One of multiple transcription start sites within the 330kb MIR100HG locus encodes a 115kb transcript that is spliced (black exons) to generate a 3kb lncRNA *(MIR100HG*) with *miR-100, miR-125b, let-7a* and the BLID protein encoded in the third intron. In CC-CR cells, *MIR100HG*, *miR-100* and *miR-125b* are highly overexpressed but *let-7a* and BLID are not (Lu et al., 2017). (B) CC-CR cells are cetuximab-resistant cells, derived from cetuximab-sensitive CC cells (Lu et al., 2017). *miR-100* and *miR-125b* are dramatically overexpressed in CC-CR cells, as measured by RT-qPCR. Data (mean±SEM, n=3) were analyzed using Students t-tests. (C) RT-qPCR analysis of *miR-100* and *miR-125b* levels in CC-CR, Δ*miR-100*, Δ*miR-125b*, Δ*miR-100/miR-125b* cell lines. Fold changes were determined using the ΔΔC(t) method. Data (mean±SEM, n=3) were analyzed using one-way Anova tests with ** indicating p<0.005, *** indicating p<0.001, and **** indicating p<0.0001.

### 3.1 Identification of mRNA Targets of *miR-100 and miR-125b*

RNAseq was performed on CC, CC-CR, and knockout cell lines (Δ*miR-100*, Δ*miR-125b*, and Δ*miR-100/miR-125b*) and bioinformatic analyses were performed to compare gene expression patterns between the lines (**Table S2**). Potential mRNA targets for *miR-100* and *miR-125b* were identified using a three-step strategy (**Fig. 2A).** First, we identified upregulated mRNAs in both CC and Δ*miR-100/miR-125b* cells compared to CC-CR cells (Lu et al., 2017). Second, we compiled a list of predicted targets for *miR-100* and *miR-125b* using three different prediction algorithms (TargetScan, TargetMiner, and miRDB; https://www.targetscan.org/vert_80/; https://www.isical.ac.in/~bioinfo_miu/targetminer20.htm; https://mirdb.org/). Third, we performed RNA immunoprecipitation of AGO2 followed by bulk and short RNA sequencing (**Tables S3,4**). AGO2 is a component of the RNA Induced Silencing Complex (RISC) in which miRNAs pair with their target mRNAs (Müller et al., 2020, Cenik and Zamore, 2011, Farazi et al., 2008, Liu et al., 2004). Immunoprecipitation of AGO2-containing complexes from CC-CR cells allowed for enrichment of *miR-100* and *miR-125b* targets because these miRNAs are so highly expressed in CC-CR cells (Lu et al., 2017). By comparing the three sets of data, we were able to identify targets of *miR-100* and *miR-125b* (**Fig. 2A**) with 51 potential targets for *miR-100* and 45 potential targets for *miR-125b* (**Fig. 2B**). When we subjected all 96 potential targets to Gene Ontology analysis using ShinyGo (Ge et al., 2020), we found significant enrichment for genes involved in cell migration, cell motility, actin filament-based processes, and MAPK signaling (**Fig. 2C; Table S5**).

**Figure 2.**
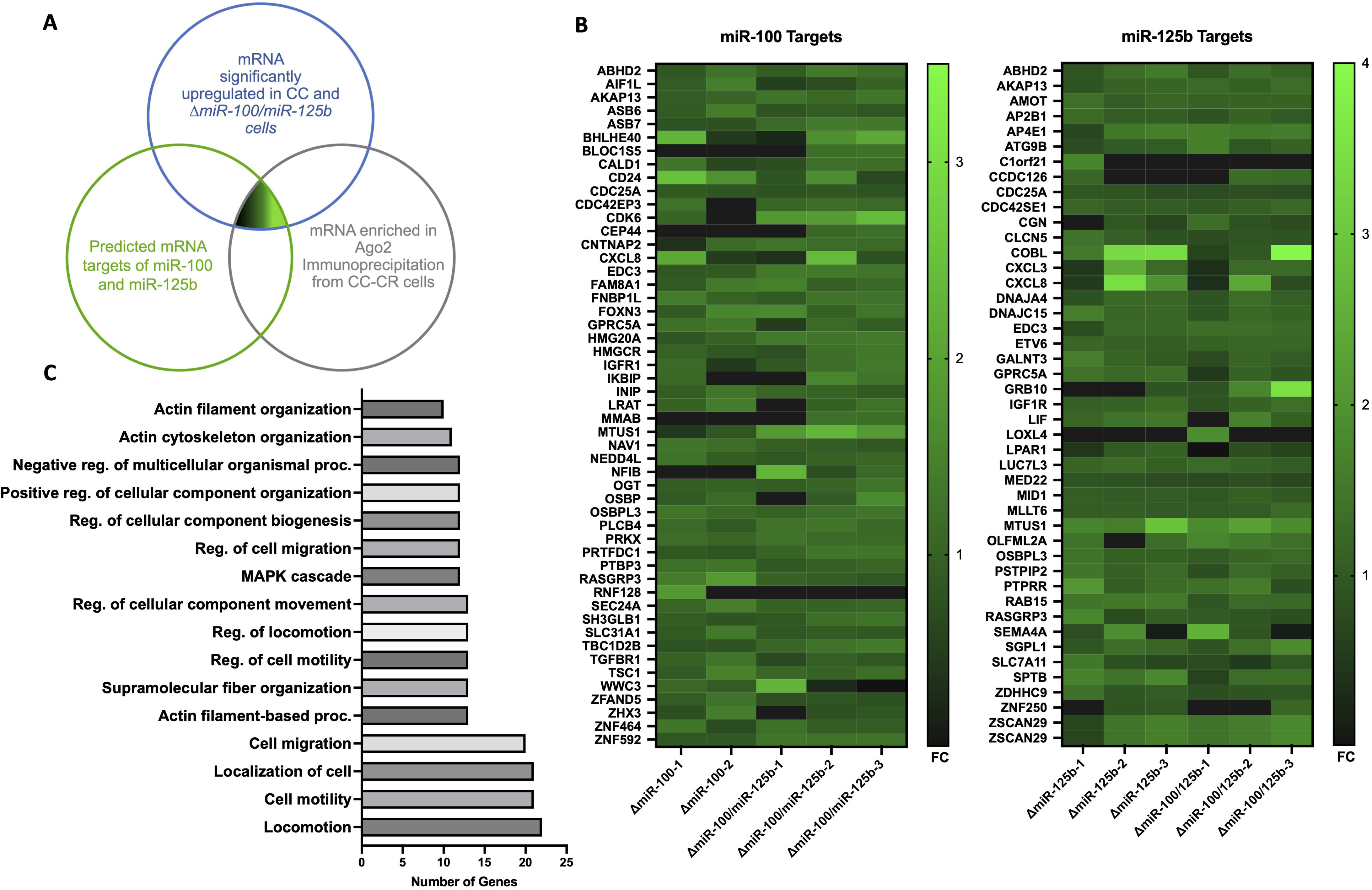
Identification of targets for *miR-100* and *miR-125b* in colorectal cancer cells. (A) Differentially expressed genes were identified by mining RNAseq data comparing CC, CC-CR, Δ*miR-100*, Δ*miR-125b*, Δ*miR-100/miR-125b* cells. Venn diagram showing overlap between genes upregulated in CC and knockout cell lines, predicted mRNA targets of *miR-100* and *miR-125b* using three different algorithms, and association with AGO2 in CC-CR cells. (B) Heatmaps were generated from the differential RNAseq data to identify potential targets upregulated in CC cells (>2 fold) and knockout cells (>1.5 fold) compared to CC-CR cells. (C) Gene Ontology analysis of the targets of *miR-100* and *miR-125b* show enrichment in genes associated with cell migration and motility. Graph was created using ShinyGO (Ge et al., 2020).

### 3.2 Silencing of mRNA Targets of *miR-100* and *miR-125b*

From the targets generated above, we examined miRNA:mRNA pairing (**Fig. S2**) and selected 15 candidate mRNAs to test for the ability of *miR-100* (3 candidates), *miR-125b* (7 candidates) or both (5 candidates) to silence luciferase reporter constructs. We designed three different categories of luciferase vectors in which the firefly luciferase open reading frame was fused to (1) no 3’-UTR (empty vector), (2) 3’-UTRs containing three perfect binding sites for *miR-100* or *miR-125b* (100/125 vector), or (3) 3’-UTRs from candidate targets (experimental vectors)(**Fig. 3A**). These vectors were transfected into either CC-CR or Δ*miR-100/miR-125b* cells and luciferase levels were determined. *Bona fide* targets of *miR-100 or miR-125b* should show increased luciferase levels in Δ*miR-100/miR-125b* cells as compared to CC-CR cells. Of the 15 targets we tested, 14 showed increased luciferase levels in Δ*miR-100/miR-125b* cells with 8 being statistically significant (**Fig. 3B**). Of the 8, CGN and PTPRR showed the most significant increase of luciferase activity when comparing Δ*miR-100/miR-125b* and CC-CR cells (**Fig. 3B**). The fact that the two most significant genes are targets of *miR-125b* is consistent with more dramatic overexpression of *miR-125b* in CC-CR cells as compared to *miR-100* (**Fig. 1B**).

**Figure 3.**
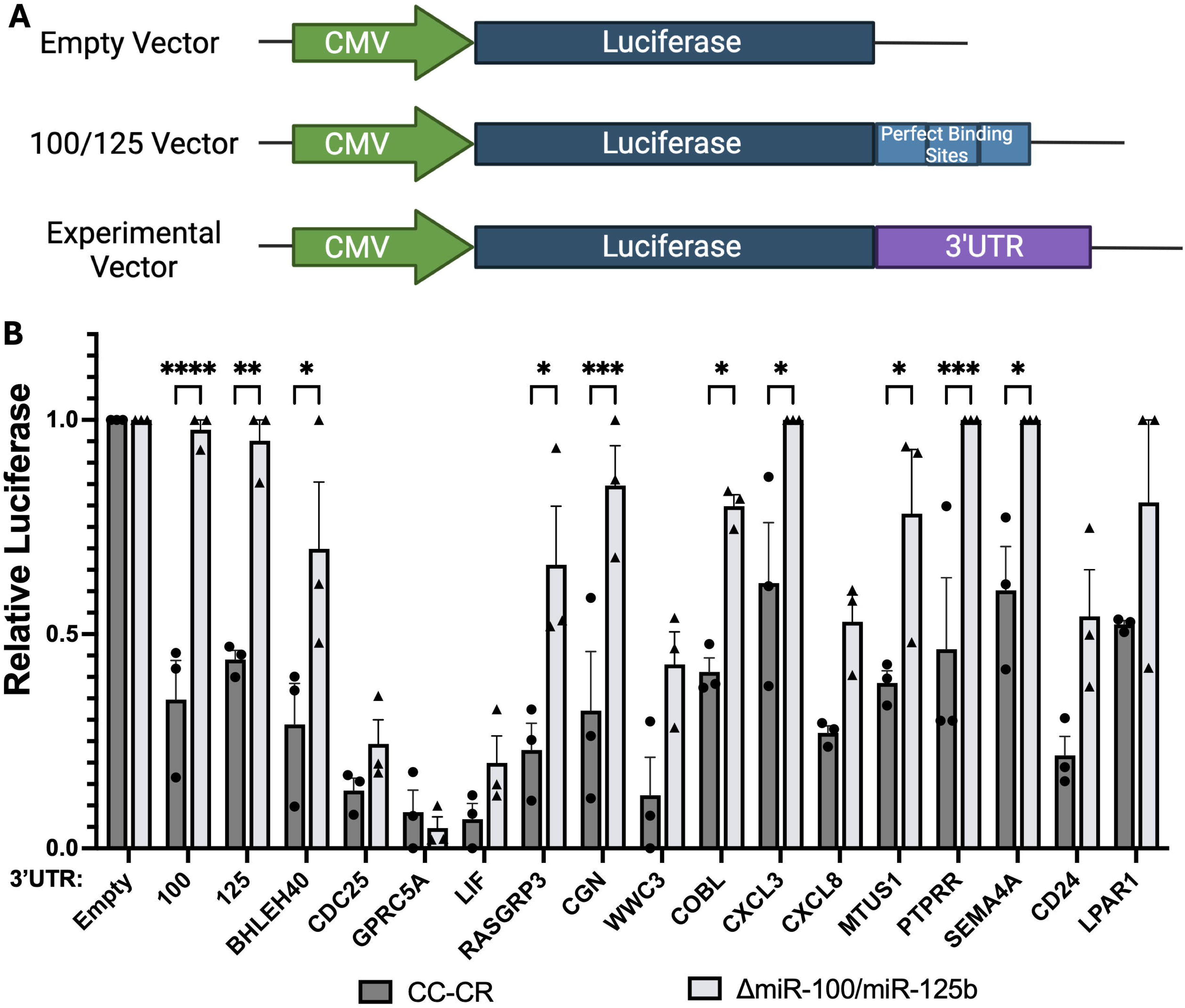
Silencing of reporter constructs containing target 3’-UTRs for *miR-100* and *miR-125b*. (A) Reporter constructs were generated in which the firefly luciferase open reading frame was fused to either plasmid 3’-UTR sequences (empty vector), 3’-UTR sequences containing three perfect binding sites for *miR-100 or miR-125b* (100/125 vector), or 3’-UTR sequences from candidate mRNA targets of *miR-100 or miR-125b* (experimental vector). (B) Quantification of relative luciferase levels after transfection of the indicated constructs into CC-CR cells or Δ*miR-100/miR-125b* cells. Relative firefly luciferase levels were calculated by normalizing to a co-transfected internal control vector expressing Renilla luciferase. Data (mean±SEM, n=3) were analyzed using two-way Anova tests with * indicating p<0.05, ** indicating p< 0.01, *** indicating p<0.001, and **** indicating p<0.0001.

### 3.3 CGN and PTPRR are Downregulated in Colorectal Cancer

*miR-100* and *miR-125b* display elevated gene expression and target multiple mRNAs in CRC and other cancer types (Li et al., 2015, Wang et al., 2020, Ottaviani et al., 2018, Cha et al., 2015, Lu et al., 2017, Zhang et al., 2021, Vallacchi and Rodolfo, 2018). CC cells grown in 3D in Type I collagen form hollow cysts with a central lumen lined by polarized cells whereas CC-CR cells form solid, disorganized colonies, indicative of aggressive behavior (Lu et al., 2017). We thus sought to test if decreased expression of CGN and PTPRR due to increased expression of *miR-100* and *miR-125b* might be observed in CRC and other cancers. We used the GENT2 database to compare the levels of CGN and PTPRR between normal and cancer tissues (**Fig. 4A,B**) and found that CGN and PTPRR are both downregulated across a variety of cancers. There are likely additional mechanisms that can lead to decreased levels of CGN and PTPRR because not all cancers show increased expression of *miR-100* (Li et al., 2015) or *miR-125b* (Wang et al., 2020), but the data are in agreement that decreased levels of CGN and PTPRR correlate with cancer.

**Figure 4.**
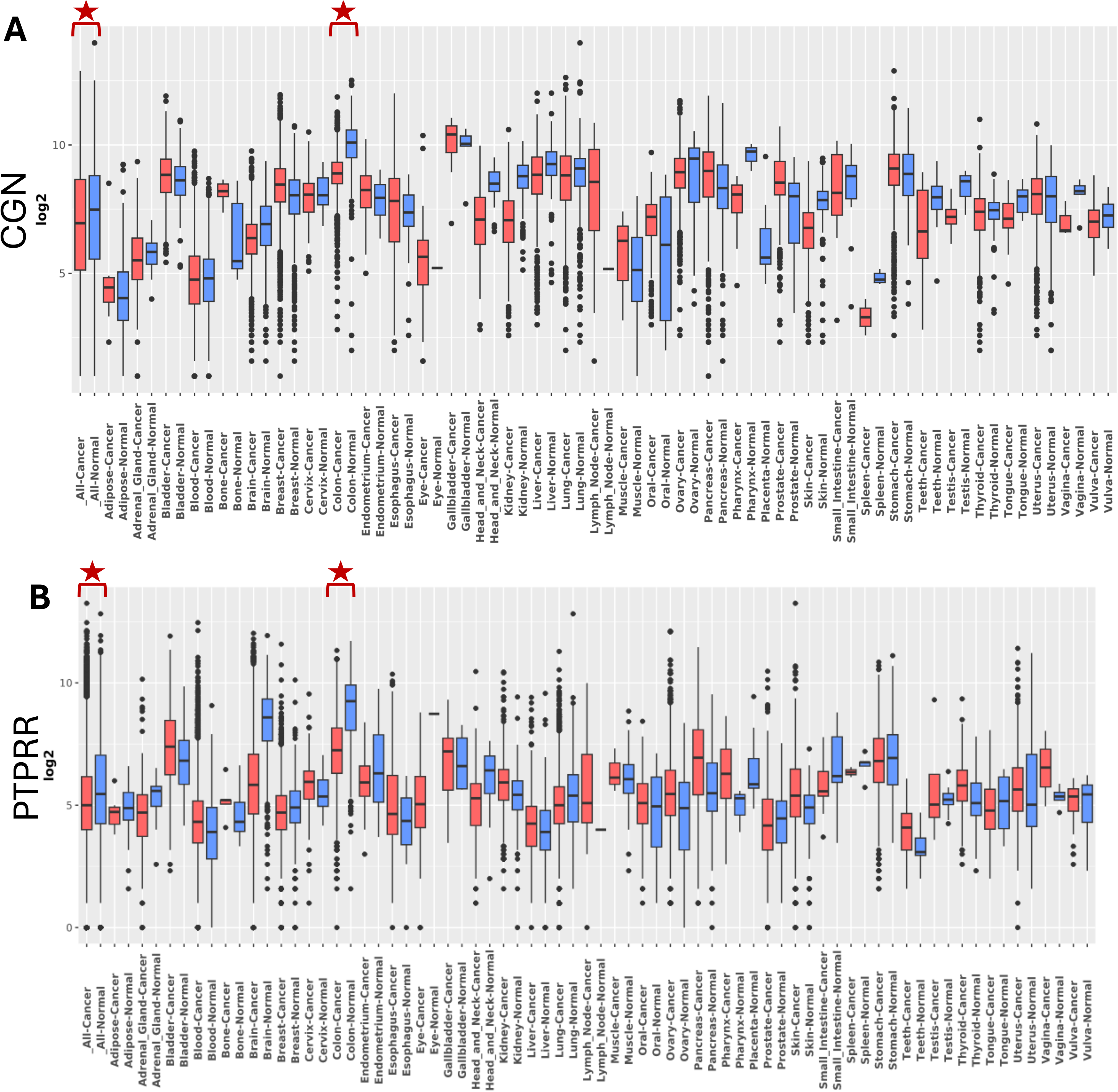
Decreased Expression of CGN and PTPRR in cancer. (A) Boxplot of CGN (A) and PTPRR (B) expression across 72 paired tissues. Red indicates cancer samples, blue indicates normal samples.

To test whether our CRC cell lines match the GENT2 data, we used antibodies to perform western blots and immunostaining of CGN protein levels in CC, CC-CR and knockout cell lines. Even though we observed silencing of PTPRR reporter constructs (Fig. 3 and see Fig. 8), we did not detect reduced protein levels by western blots, so we decided to focus hereafter on CGN. CGN connects tight junctions to the cytoskeleton and localizes to apical junctions joining adjacent epithelial cells (Zhang et al., 2021, Citi et al., 1988). In cells that express low to undetectable levels of *miR-100 or miR-125b* (CC and Δ*miR-100/miR-125b* cells), CGN was readily detected by western blots (**Fig. 5A,B**) and localized to cell-cell junctions by immunofluorescence (**Fig. 5C**). Immunofluorescent signals were more readily visualized in CC cells compared to Δ*miR-100/miR-125b* cells, potentially due to regulation of other factors by *miR-100 or miR-125b* that also seem to affect overall morphology, as indicated by differences in F-actin staining and changes in CGN localization. In cells expressing high levels of *miR-100* and *miR-125b* (CC-CR cells), CGN detection by both western blots and immunofluorescence was dramatically reduced (**Fig. 5**). These results are consistent with targeting of CGN by *miR-100* and *miR-125b*.

**Figure 5.**
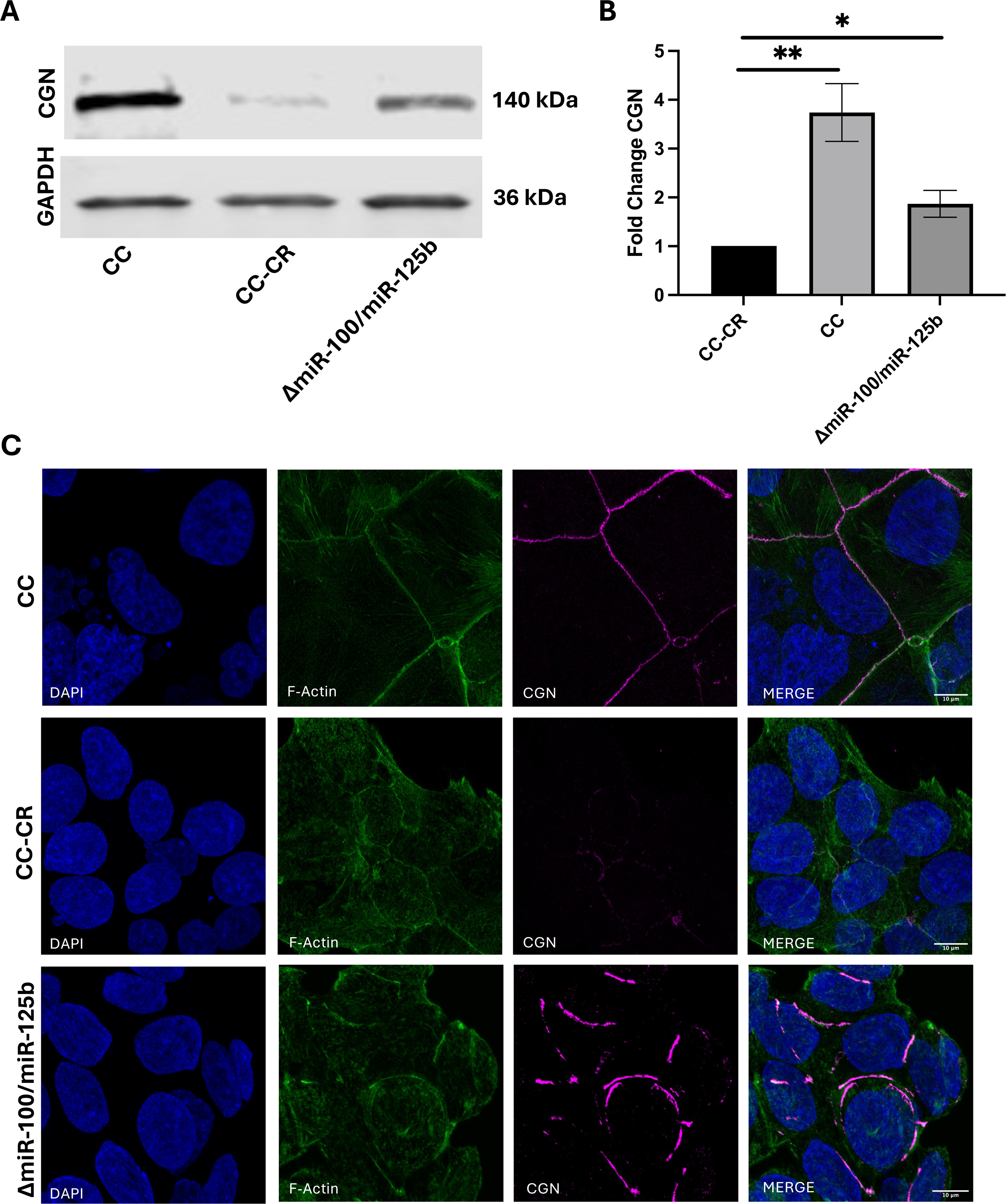
Decreased Expression of CGN in Cells Expressing High Levels of *miR-100* and *miR-125b*. (A,B) Western blots and quantitation of CGN levels in the indicated cell lines. (C) The indicated cells were fixed and immunostained with DAPI to stain nuclei (blue), antibodies against F-actin (green), or antibodies against CGN (magenta). Merged images are shown at the far right. Scale bar indicates 10μm. For B, data (mean±SEM, n=3) were analyzed using Students t-tests. For significance, *indicates p<0.05, **indicates p< 0.01.

### 3.4 *miR-100* and *miR-125b* Contribute to Enhanced 3D Growth and Invasiveness in CRC cells

Gene Ontology analysis of targets of *miR-100* and *miR-125b* showed enrichment of genes associated with cell migration and motility (**Fig. 2C**). This is consistent with downregulation of CGN which leads to increased invasiveness, metastatic potential, and EMT (Guillemot and Citi, 2006, Mangan et al., 2016, Schossleitner et al., 2016, Zhang et al., 2021). To test the effect of *miR-100* and *miR-125b* and their targeting of CGN on 3D growth, we cultured CC, CC-CR and Δ*miR-100/miR-125b* cells under spheroid growth conditions (Carey et al., 2013) in low adherence plates for 3 days before transfer to growth in Type I collagen. Under these conditions at both day 0 and day 5 in collagen, the morphology of Δ*miR-100/miR-125b* colonies was more similar to CC colonies than to colonies derived from parental CC-CR cells (**Fig. 6A**; **Supp. Movies 1-3**). At day 5, we also observed significant differences in surface dynamics when comparing colonies derived from the three lines **(Supp. Movies 1-3)**. Increased projections emanating from the edges of the colonies and protruding into collagen were observed with CC-CR cells, whereas far fewer projections were observed in colonies derived from CC cells grown under the same conditions. As with overall morphology, Δ*miR-100/miR-125b* colony dynamics and projections were more similar to that observed in CC colonies than CC-CR colonies **(Supp. Movies 1-3).**

**Figure 6.**
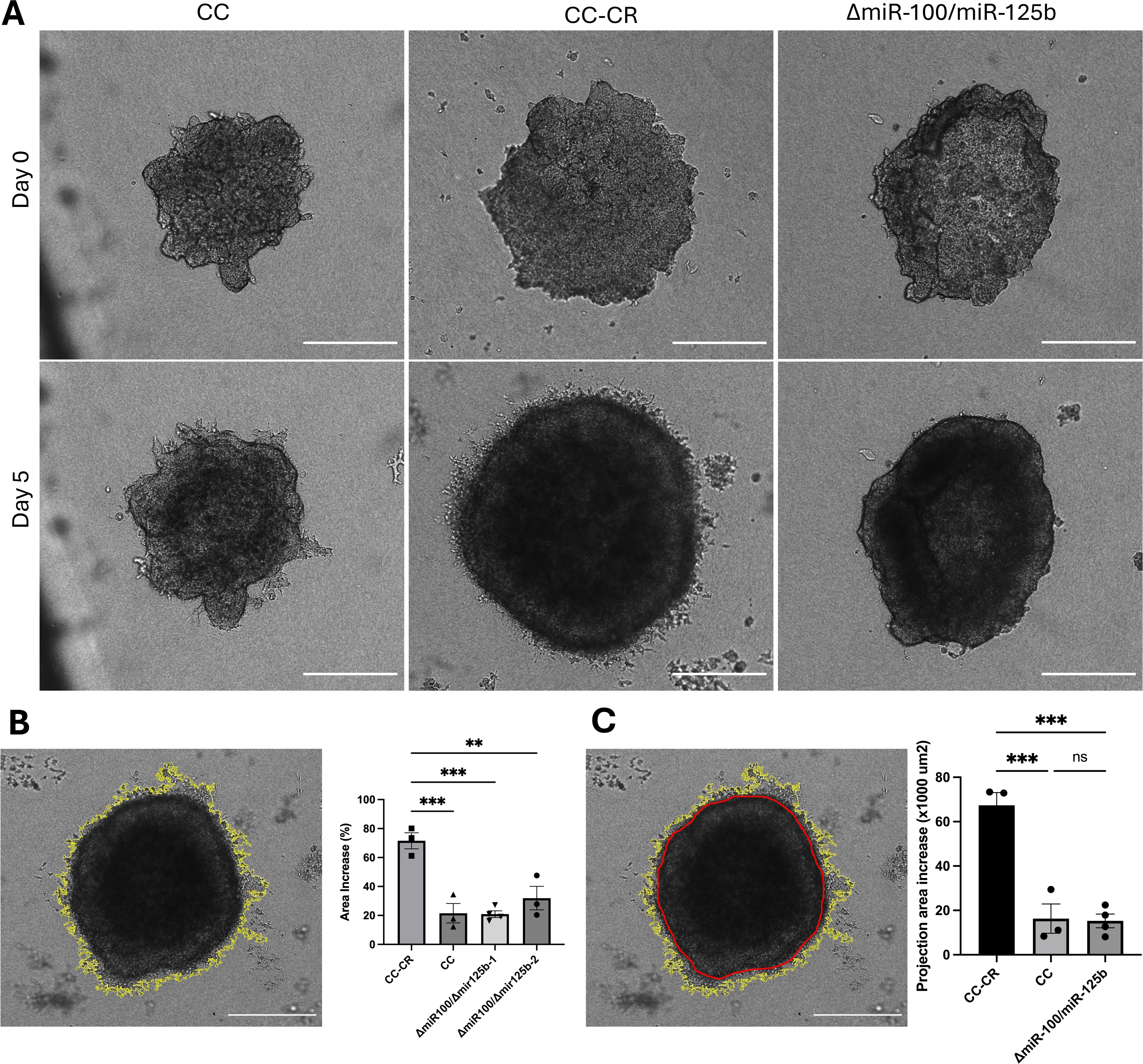
*miR-100* and *miR-125b* contribute to enhanced 3D growth and invasiveness. (A) The indicated cell types were plated in low adherent, spheroid cultures for 3 days before transfer to growth in Type I collagen and live imaging over 5 days. (B) The increase in colony area was determined by measuring the surface area of spheroids from day 0 to day 5. Surface area tracing was performed using ImageJ, shown in yellow. Significance was analyzed using one-way ANOVA tests, (n=3). (C) Increases in area due to colony projections into the collagen (invasiveness) were calculated by measuring the total surface area including projections (yellow) and subtracting the surface area of the main spheroid (red). Significance was analyzed using one-way ANOVA tests, data (n=3). Data represent mean ± SEM with **indicating p< 0.01, ***indicating p<0.001; ns=not significant.

Quantification of overall spheroid growth revealed that colonies derived from CC-CR cells showed a significant increase in total spheroid area when compared to colonies derived from CC or Δ*miR-100/miR-125b* cells (**Fig. 6B**). We also quantified the number and size of projections into the collagen as a measure of invasiveness by determining the increase in projection surface area after growth in collagen for five days. CC-CR colonies showed a significantly greater increase in projection area compared to CC and Δ*miR-100/miR-125b* colonies (**Fig. 6C**). While invasiveness is likely due to multiple changes, the decreased projections we observe in CC and Δ*miR-100/miR-125b* colonies are consistent with increased expression of CGN due to lower levels of *miR-100* and *miR-125b*.

### 3.5 *miR-100* and *miR-125b* Contribute to Enhanced 3D Growth and Invasiveness in Glioblastoma cells

To test whether targeting of CGN by *miR-100* and *miR-125b* can alter the growth characteristics of non-CRC cells, we used the LN-18 glioblastoma line (Diserens et al., 1981). To our surprise, these cells express even higher levels of *miR-100* and *miR-125b* than CC-CR cells, consistent with undetectable levels of CGN (**Fig. 7A-D**), consistent with the GENT2 analysis in **Fig. 4A**. When we plated LN-18 cells under spheroid conditions and transferred the colonies to growth in 3D type I collagen over 5 days, we observed an increase in colony growth (**Fig. 7E**). However, when we overexpressed CGN levels in LN-18 cells, we detected a significant decrease in colony growth (**Fig. 7E**). This demonstrates that CGN levels can affect colony growth and that the effect of CGN is not restricted to CRC cells.

**Figure 7.**
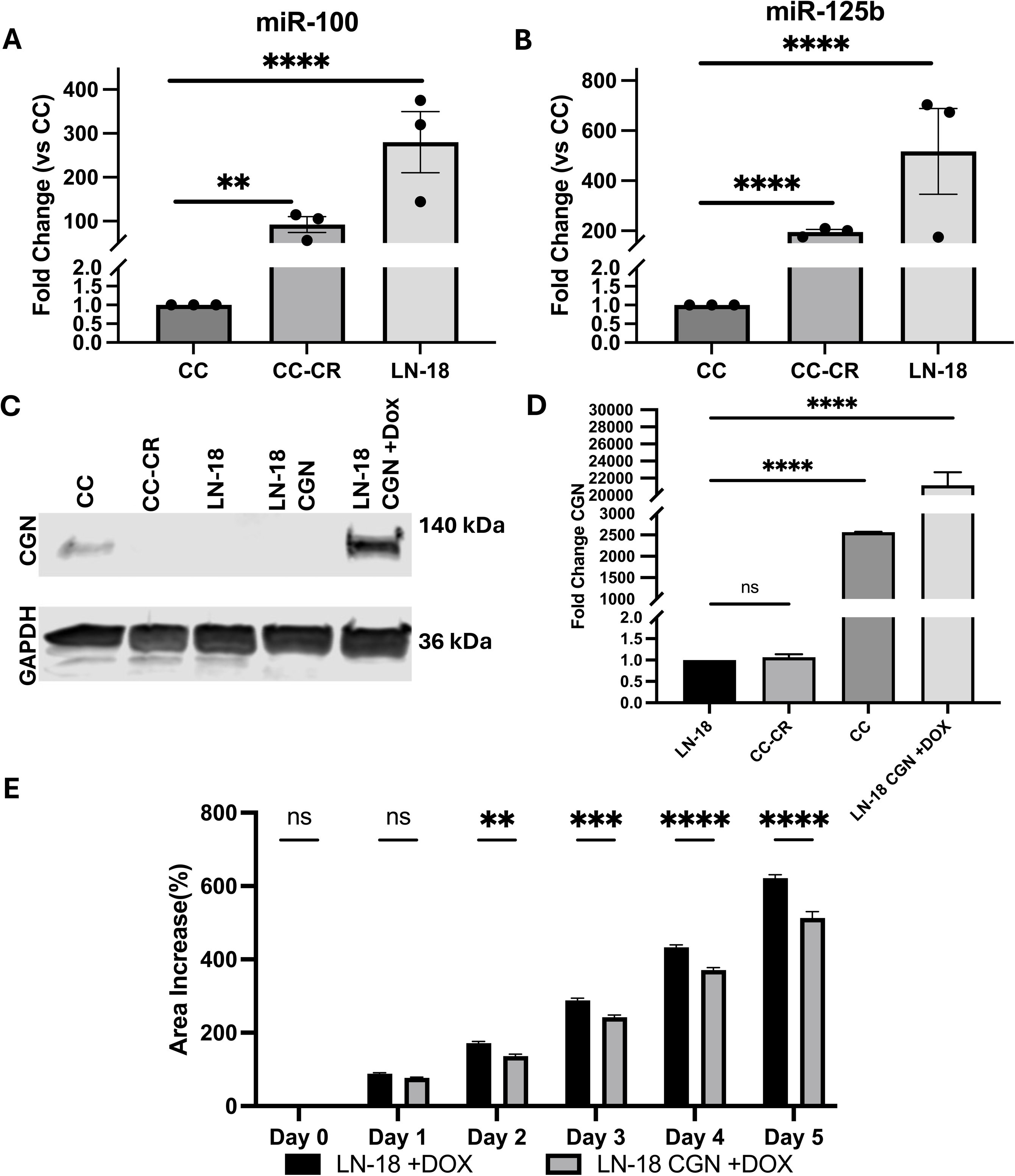
Increased CGN expression in glioblastoma contributes to decreased 3-D growth. (A,B) Relative levels of *miR-100* and *miR-125b* in CC, CC-CR, and LN-18 (glioblastoma) cell lines. Data represent mean ± SEM with **indicating p< 0.01, and **** indicating p<0.0001. Significance was determined using Students t-tests; n=3. (C) Cingulin (CGN) levels were determined using western blots in CC, CC-CR, LN-18 and doxycyclin inducible LN-18 cell lines. Lane loading was monitored with blots against GAPDH. (D) Quantification of blots as in C showing fold-changes in CGN in the indicated cell lines normalized to GAPDH. Data represent mean ± SEM with n=3. Significance was determined using Students t-tests with **** indicating p<0.0001; ns=not significant. (E) Increase in spheroid area was measured after transfer to collagen from 0-5 days. Surface area tracing was performed using ImageJ. Significance was determined using two-way ANOVA tests, n=6. Data represent mean ± SEM with *indicating p<0.05, **indicating p< 0.01, ***indicating p<0.001, **** indicating p<0.0001; ns=not significant.

### 3.6 Extracellular Transfer of *miR-100* and *miR-125b*

Previous work identified CGN and other targets for *miR-100* and *miR-125b* and showed that cellular changes in these targets can drive migration and invasion of CRC cells (Zhang et al., 2021). Here, we sought to extend those experiments to focus on extracellular transfer of *miR-100* and *miR-125b* because we had previously shown that these miRNAs can be secreted from CRC cells (Cha et al., 2015, Abner et al., 2021). However, definitive evidence of extracellular transfer of these miRNAs was lacking because the experiments used recipient cells that express *miR-100* and *miR-125b* which can lead to indirect silencing of reporter constructs and potential false positives (Gruner and McManus, 2021). Our knockout cell lines provide an ideal experimental system to conclusively test for extracellular transfer of *miR-100* and *miR-125b* between donor and recipient cells. As donor cells, we used CC-CR cells because they not only express high levels of *miR-100* and *miR-125b*, but they also secrete both miRNAs (**Fig. 8A).** Thus, we set up Transwell assays with CC-CR donor cells and Δ*miR-100/miR-125b* recipient cells and co-cultured the cells for 5 days before isolation of RNA from the recipient cells and analysis of *miR-100* and *miR-125b* levels. Compared to control transfer experiments in which Δ*miR-100/miR-125b* cells were used as both the donor and recipient cells, we observed dramatic increases in the levels of *miR-100* and *miR-125b* in Δ*miR-100/miR-125b* recipient cells (∼150- and 60-fold, respectively) when CC-CR cells were used as the donor (**Fig. 8B**).

**Figure 8.**
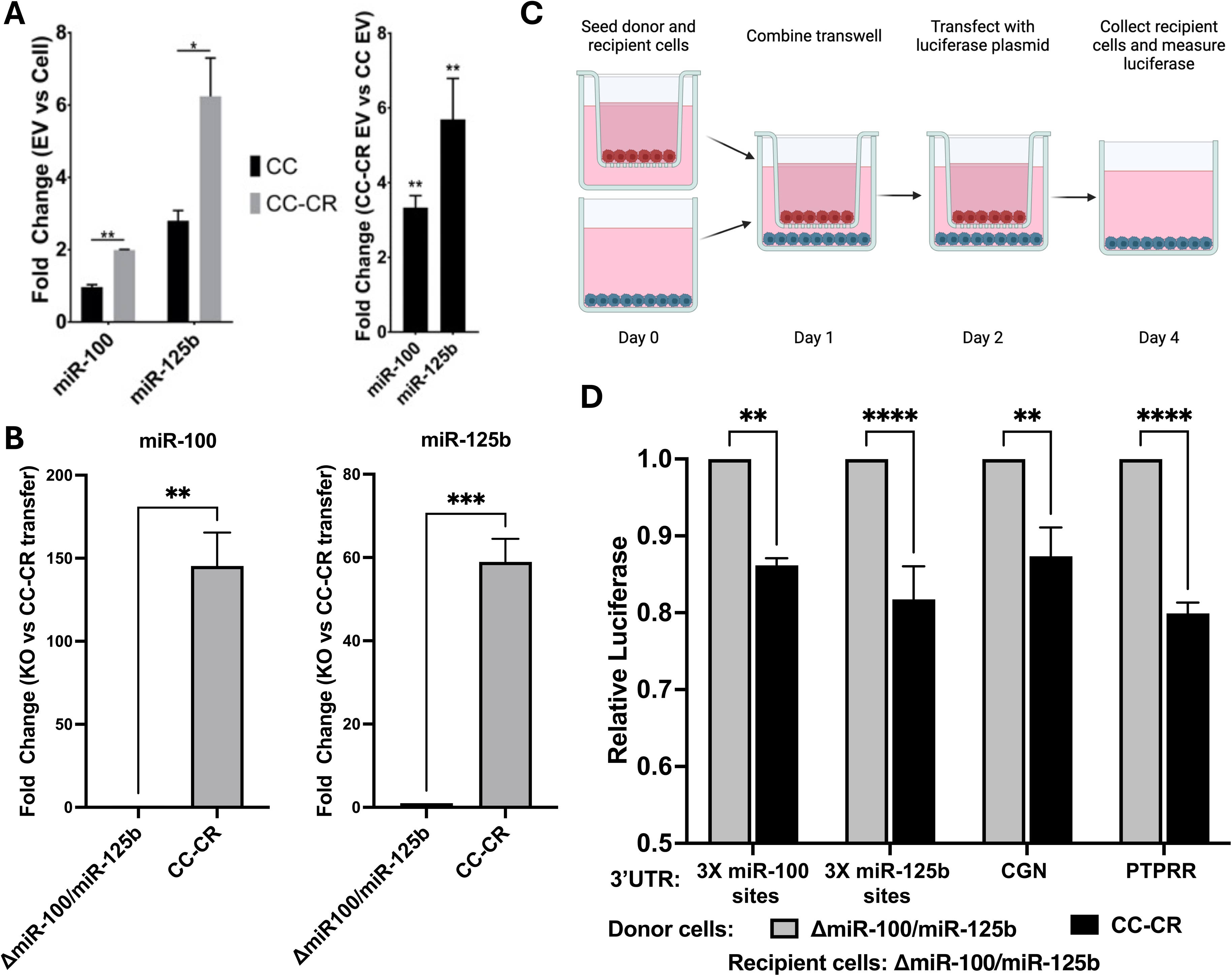
Functional transfer of *miR-100* and *miR-125b*. (A) RNA was collected from extracellular vesicles (EVs) from CC and CC-CR cells and the levels of *miR-100* and *miR-125b* were quantified by RT-qPCR. The left panel shows the fold-change enrichment of these miRNAs in EVs compared to cellular levels. The right panel shows the fold-change in EV levels of *miR-100* and *miR-125b* when comparing CC-CR and CC cells. (B) Transwell co-culture experiments were performed with Δ*miR-100/miR-125b* recipient cells and the donor cells as indicated on the X axis. RNA was collected from the recipient cells after 5 days in culture and fold-changes in *miR-100* and *miR-125b* levels were determined by RT-qPCR. (C) Schematic of Transwell co-culture experiments to test silencing of CGN and PTPRR luciferase reporter constructs in Δ*miR-100/miR125b* recipient cells. (D) Δ*miR-100/miR-125b* recipient cells were transfected with luciferase reporters and co-cultured with either Δ*miR-100/miR-125b* (grey) or CC-CR (black) donor cells. Relative luciferase values (y-axis) were calculated in recipient cells, normalizing as in Fig. 3. For A and B, data (mean±SEM, n=3) were analyzed using one-way Anova tests while the data in panel D (mean±SEM, n=3) were analyzed using two-way Anova tests. For significance, *indicates p<0.05, **indicates p< 0.01, ***indicates p<0.001, and **** indicates p<0.0001; ns is not significant.

Knowing that *miR-100* and *miR-125b* can be successfully transferred between cells, we tested whether their transfer from CC-CR cells to Δ*miR-100/miR-125b* recipient cells can result in functional miRNA transfer. For this, we used Transwell co-culture assays in which the Δ*miR-100/miR-125b* recipient cells were transfected with vectors in which the firefly luciferase open reading frame was fused to the 3’-UTRs from either CGN or PTPRR (**Fig. 8C**). Compared to controls, we observed a significant reduction in luciferase levels when CC-CR cells were the donor cells and Δ*miR-100/miR-125b* were the recipient cells (**Fig. 8D**). These data demonstrate that *miR-100 and miR-125b* can be functionally transferred between cells to silence reporter mRNAs. To test whether transfer of these miRNAs can silence endogenous CGN in recipient cells, we used Transwell co-culture experiments with Δ*miR-100/miR-125b* recipient cells and either CC-CR or Δ*miR-100/miR-125b* donor cells followed by immunofluorescent staining of CGN in recipient cells. With Δ*miR-100/miR-125b* donor cells, CGN immunostaining was readily apparent in Δ*miR-100/miR-125b* recipient cells (**Fig. 9A**) at levels similar to those observed in **Fig. 5C**. However, when CC-CR cells were used as the donor cells, CGN was undetectable in Δ*miR-100/miR-125b* recipient cells (**Fig. 9A**). These results demonstrate that endogenous CGN can be targeted by functional transfer of *miR-100* and *miR-125b* leading to decreased endogenous target protein levels.

**Figure 9.**
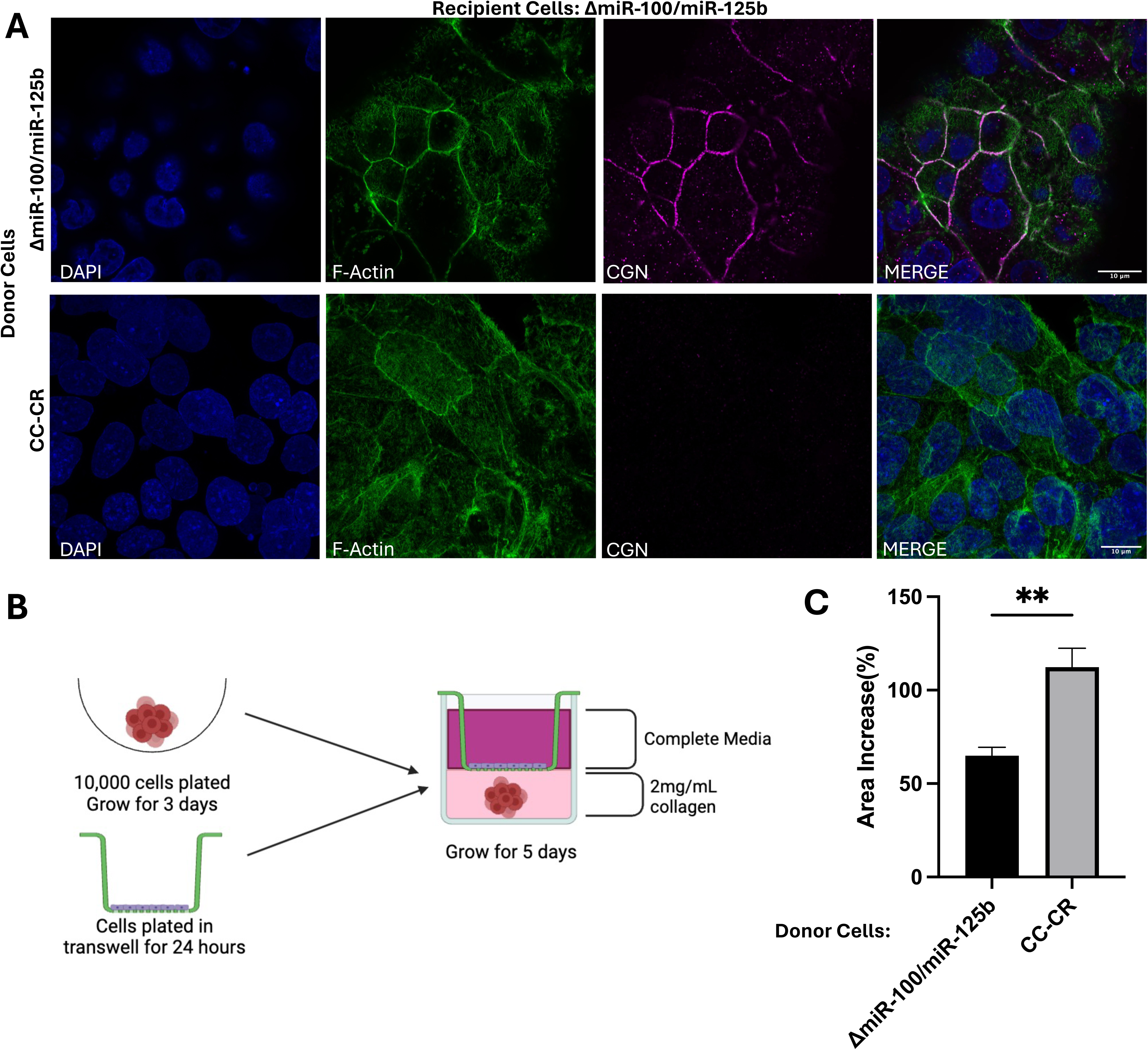
Transfer of *miR-100* and *miR-125b* decreases CGN immunofluorescence in recipient cells. (A) Transwell co-culture experiments were performed with Δ*miR-100/miR-125b* recipient cells and the indicated donor cells. After 5 days, recipient cells were immunostained as in Fig. 5C. Scale bar indicates 10μm. B. Schematic of Transwell co-culture experiments to test growth changes in spheroids due to transfer of material from donor 2-D cells to recipient spheroids grown in 3D. (C) The increase in spheroid colony area was determined by measuring the surface area of spheroids from day 0 to day 5 after exposure to donor cells. Significance was determined using Students t-test; n=6. Data represent mean ± SEM with **indicating p< 0.01.

As a final assay to test for functional extracellular transfer of *miR-100* and *miR-125b*, we used Transwell experiments in which spheroid cultures were plated in 3D type I collagen and co-cultured with either CC-CR or Δ*miR-100/miR-125b* cells. Exposure of spheroid colonies to CC-CR cells which express high levels of *miR-100* and *miR-125b* led to an increase in the area of the resultant colonies when compared to co-culture with Δ*miR-100/miR-125b* cells (**Fig. 9B**). This supports the hypothesis that extracellular transfer of *miR-100* and *miR-125b* can alter 3D growth characteristics and invasiveness, consistent with reduced levels of CGN.

## 4. Discussion

While many reports have suggested that miRNAs can undergo extracellular transfer between cells (Mittelbrunn et al., 2011, Valadi et al., 2007, Skog et al., 2008, Boelens et al., 2014, Squadrito et al., 2014, Cha et al., 2015, Barman et al., 2022), the recipient cells in nearly all of these experiments express the same miRNAs, complicating interpretation of target mRNA silencing after transfer (Gruner and McManus, 2021). Silencing under these conditions could be due to indirect effects including unexpected increased expression of endogenous miRNAs. Our knockout cell lines provide an ideal experimental system to obviate such concerns. Using donor cells that express high levels of *miR-100* and *miR-125b* and recipient cells with targeted deletions of these genes, we identified mRNA targets and show functional transfer of *miR-100* and *miR-125b* to silence both reporter constructs and endogenous genes in recipient cells.

All cells release a heterogeneous mixture of membrane-bound vesicles and nanoparticles with current work devoted to understanding biogenesis and cargo loading mechanisms (Dixson et al., 2023). Our experiments conclusively demonstrate extracellular trafficking of *miR-100* and *miR-125b* but whether they are transferred as RNA-protein complexes, nanoparticles (exomeres and supermeres), or membrane-encased small or large vesicles remains to be determined (Jeppesen et al., 2023, Zhang et al., 2019).

### 4.1 Tumor Microenvironment Implications

Invasiveness and metastasis are indicative of differing stages of cancer and can also be indicative of patient prognosis (Hanahan and Weinberg, 2011, Zeineddine et al., 2023). EMT is a pivotal stage of metastasis involving the breakdown of cell-cell adhesions (Pretzsch et al., 2019). Located on apical regions of adjacent epithelial cells, tight junctions play a crucial role in cell-cell adhesion and tissue integrity. Disruption of tight junctions can allow cells to become invasive and metastatic (Martin, 2014). CGN serves as a tether between the cytoskeleton and tight junction components, interacting with ZO-1 at apical junctions (Zihni et al., 2016, Citi et al., 1988). Loss of CGN in epithelial cells disrupts tight junction formation and cell polarity (Mangan et al., 2016, Guillemot and Citi, 2006, Schossleitner et al., 2016). Coupled to our finding that *miR-100* and *miR-125b* are enriched in EVs from CRC cells and that *miR-100* and *miR-125b* can be transferred between cells, our data support a model whereby cell-cell communication within the tumor microenvironment can alter tight junctions leading to altered growth and increased metastatic potential in recipient cells.

Disruption of tight junctions can also alter the blood-brain barrier in glioblastoma, referred to as the blood-tumor barrier (Steeg, 2021). In this case, tight junctions and adherens junctions in endothelial cells are disrupted leading to impaired barrier function, development of EMT, and increased invasiveness (Iwadate, 2016). Our finding that glioblastoma cells express high levels of *miR-100* and *miR-125b* and correspondingly low levels of CGN are consistent with the increased invasiveness we observe in 3D colony growth and further support the finding that disruption of tight junctions through decreased CGN levels can facilitate tumor progression through both cell-autonomous and non-cell-autonomous mechanisms.

Our experiments focus on cell-cell communication within the tumor microenvironment using Transwell assays with donor and recipient cells. It should be emphasized that such communication can be extensive, with both tumor and normal (stromal) cells secreting vesicles and nanoparticles that can engage in two-way exchange. The exchange of miRNAs and other cargo within the tumor environment can not only increase tumor growth, invasiveness and metastasis, but also modulate anti-tumor immune responses (Reale et al., 2022).

### 4.2 MIR100HG, Tumor Growth, Invasiveness and EMT

Beyond demonstrating miRNA transfer between cells, our knockout lines provided a model to test the effects of 3D growth in the presence and absence of *miR-100* and *miR-125b.* In 3D growth in collagen, CC cells form hollow cysts with a central lumen lined by polarized cells whereas CC-CR cells grow into the central lumen with disorganized, solid colonies (Lu et al., 2017). A prior deletion of exon 4 within the MIR100HG locus supported the idea that MIR100HG contributes to both cetuximab resistance and metastasis (Liu et al., 2022). Our precise deletions of *miR-100* and *miR-125b* allowed us to test the effect of these miRNAs on 3D growth and showed that loss of these miRNAs results in slower overall growth with decreased edge dynamics and fewer invasive projections. Since expression of MIR100HG is unaffected in our knockout lines, our data agree with previous results that *MIR100HG* can induce EMT, but also that *miR-100* and *miR-125b* play a similar role through downregulation of CGN (Zhang et al., 2021). The data are also consistent with our GO analysis of the targets of *miR-100* and *miR-125b* being enriched in cell motility and migration (**Fig 2C**).

In summary, our data support findings that increased expression of *miR-100* and *miR-125b* in cancer promotes disruption of tight junctions, activation of EMT, and increased metastatic potential. Given that these miRNAs are abundant in EVs secreted from CRC and glioblastoma cells, the data support both cell-autonomous and non-cell-autonomous roles for *miR-100* and *miR-125b*.

## 5. Acknowledgements

The authors would like to thank all members of the Coffey, Weaver, and Patton labs for advice and suggestions. The authors would also like to thank the Coffey and King labs for the generous gift of the cell lines.

## 6. Funding

This work was supported by PO1 CA229123 to RJC, AMW, KCV, QL, and JGP and an American Cancer Society Research Scholar grant to MR.

## 7. Author Contributions

HMN, SQ, LH, SC, NL, SS, LT, MS, LG, and KCC generated all data in the manuscript. KCV, QL, JLF, AMW, MR, and RJC provided scientific advice and suggestions and edited the manuscript. HMN, SQ, JLF, MR and JGP wrote the paper.

## Notes

### Competing Interest Statement

The authors have declared no competing interest.

### Summary of Updates

We have extensively re-written the manuscript with additional figures and data. An additional cell line as well as clarifications have been added. All figures have been revised.

