## Supplementary figures and images for "*miR-100* and *miR-125b* Contribute to Enhanced 3D Growth and Invasiveness and can be Functionally Transferred to Silence Target Genes in Recipient Cells"

### Supplemental Figure 1

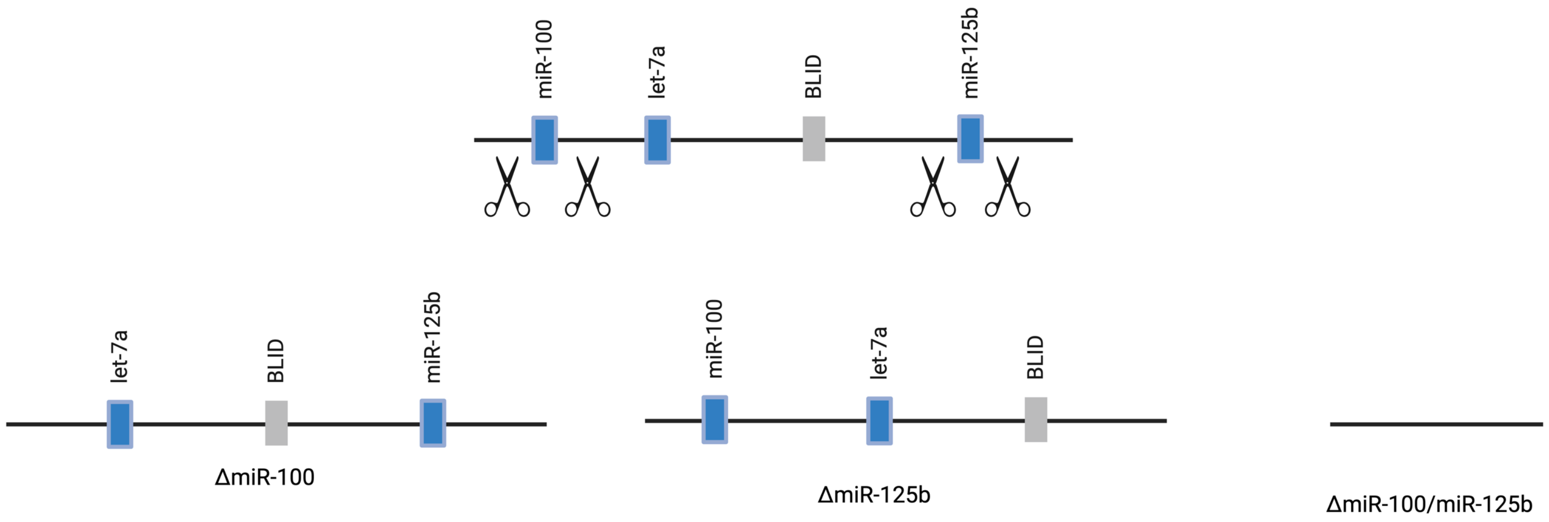

### Supplemental Figure 2

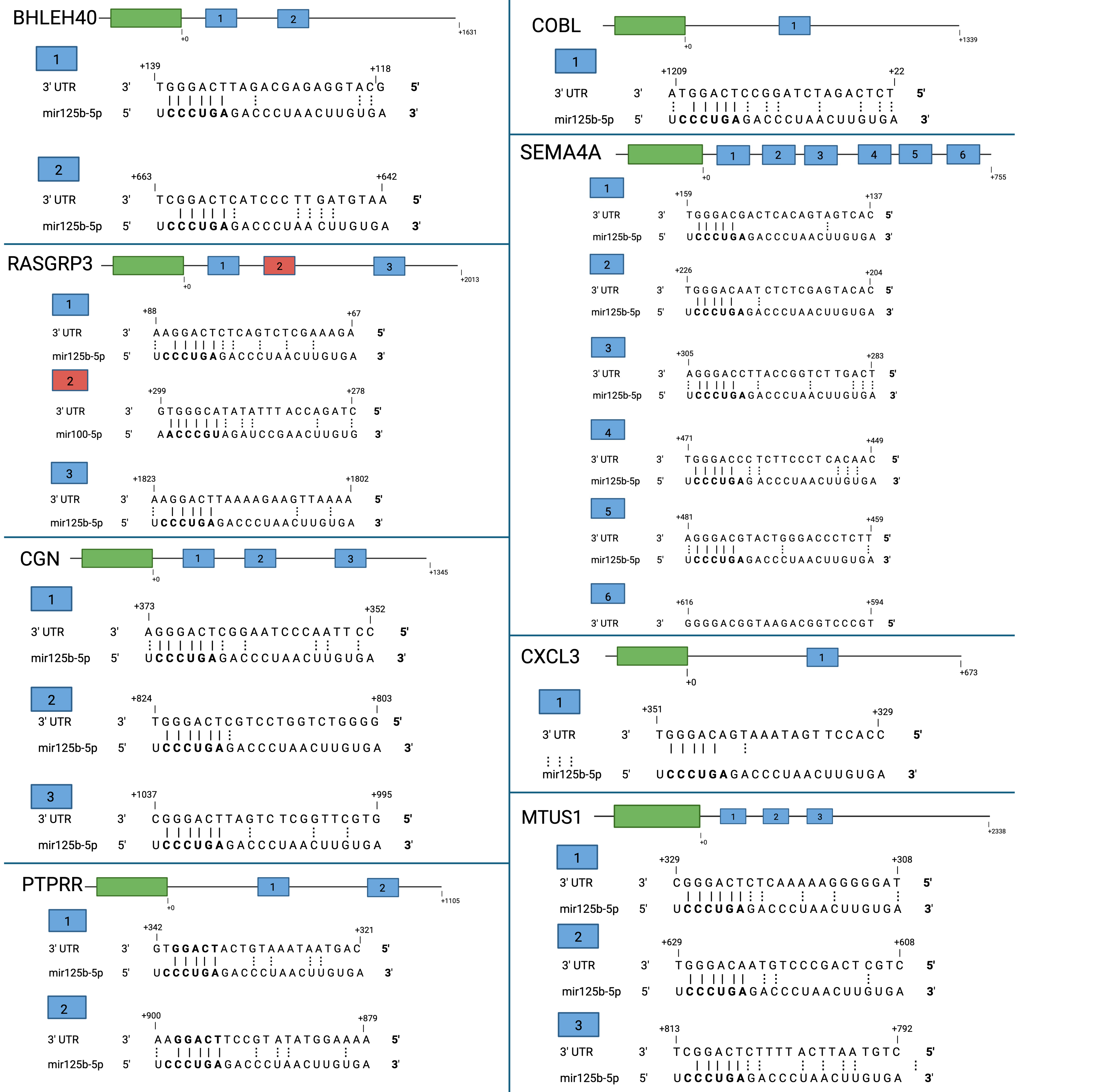
